# The Impact of In-Scanner Head Motion on Structural Connectivity Derived from Diffusion Tensor Imaging

**DOI:** 10.1101/185397

**Authors:** Graham L. Baum, David R. Roalf, Philip A. Cook, Rastko Ciric, Adon F.G. Rosen, Cedric Xia, Mark A. Elliot, Kosha Ruparel, Ragini Verma, Birkan Tunc, Ruben C. Gur, Raquel E. Gur, Danielle S. Bassett, Theodore D. Satterthwaite

**Author notes:** Please address correspondence to: Theodore D. Satterthwaite, M.D., 10th Floor, Gates Building, Hospital of the University of Pennsylvania 34th and Spruce Street Philadelphia, PA 19104.

## Abstract

Multiple studies have shown that data quality is a critical confound in the construction of brain networks derived from functional MRI. This problem is particularly relevant for studies of human brain development where important variables (such as participant age) are correlated with data quality. Nevertheless, the impact of head motion on estimates of structural connectivity derived from diffusion tractography methods remains poorly characterized. Here, we evaluated the impact of in-scanner head motion on structural connectivity using a sample of 949 participants (ages 8-23 years old) who passed a rigorous quality assessment protocol for diffusion tensor imaging (DTI) acquired as part of the Philadelphia Neurodevelopmental Cohort. Structural brain networks were constructed for each participant using both deterministic and probabilistic tractography. We hypothesized that subtle variation in head motion would systematically bias estimates of structural connectivity and confound developmental inference, as observed in previous studies of functional connectivity. Even following quality assurance and retrospective correction for head motion, eddy currents, and field distortions, in-scanner head motion significantly impacted the strength of structural connectivity in a consistency-and length-dependent manner. Specifically, increased head motion was associated with reduced estimates of structural connectivity for high-consistency network edges, which included both short-and long-range connections. In contrast, motion inflated estimates of structural connectivity for low-consistency network edges that were primarily shorter-range. Finally, we demonstrate that age-related differences in head motion can both inflate and obscure developmental inferences on structural connectivity. Taken together, these data delineate the systematic impact of head motion on structural connectivity, and provide a critical context for identifying motion-related confounds in studies of structural brain network development.

## INTRODUCTION

Diffusion tensor imaging (DTI) remains the most commonly-used technique for characterizing human white matter (WM) microstructure *in vivo* (Assaf and Pasternak, 2008; Basser et al., 1994; Basser and Pierpaoli, 1996). Graph theoretical analysis of diffusion tractography data has provided a fruitful quantitative framework for delineating how structural brain architecture shapes intrinsic functional activity and cognition (Bullmore and Sporns, 2009; Rubinov and Sporns, 2010), particularly in the context of human brain development (Baum et al., 2017; Grayson et al., 2014; Hagmann et al., 2010) and neuropsychiatric disorders (Bassett et al., 2008; Bohlken et al., 2016; Collin et al., 2017; Di Martino et al., 2014; Kessler et al., 2016; Satterthwaite et al., 2015; Sun et al., 2017). Nonetheless, prior work has shown that artifacts caused by eddy currents, head motion, and magnetic susceptibility can negatively impact diffusion model fitting and subsequent microstructural measures (Jones and Basser, 2004; Le Bihan et al., 2006).

Despite recent focus on the influence of head motion on data quality in other imaging modalities including resting state functional connectivity (Fair et al., 2012; Power et al., 2012; Satterthwaite et al., 2012; Van Dijk et al., 2012; C.-G. Yan et al., 2013) and structural imaging (Alexander-Bloch et al., 2016; Pardoe et al., 2016; Reuter et al., 2015; Savalia et al., 2017; Tisdall et al., 2012, 2016), the impact of motion on structural connectivity derived from diffusion tractography remains sparsely investigated. Prior work using DTI has demonstrated that head motion increases the uncertainty of diffusion model fitting (Bastin et al., 1998; Landman et al., 2007; Ling et al., 2012; Tijssen et al., 2009), impacting the estimation of diffusion scalar measures such as fractional anisotropy (FA) and mean diffusivity (MD). These measures are highly sensitive (but not specific) to underlying WM microstructural properties such as axonal packing density and myelination (Chang et al., 2017; Gulani et al., 2001; Takahashi et al., 2002). Notably, motion artifact can produce artificially higher FA in low anisotropy gray matter regions (Bastin et al., 1998; Farrell et al., 2007; Landman et al., 2008), while simultaneously leading to diminished FA in high anisotropy WM regions (Aksoy et al., 2008; Jones and Basser, 2004; Le Bihan et al., 2006). These spurious effects may impact streamline tractography algorithms during the step-wise reconstruction of WM pathways, particularly when streamline termination criteria are defined by local FA and angular thresholds (Girard et al., 2014).

While image processing tools have been developed to retrospectively estimate and mitigate the influence of motion artifact on diffusion-weighted images (Andersson et al., 2016; Andersson and Sotiropoulos, 2016; Rohde et al., 2004), important work by Yendiki et al. (2014) and others (Liu et al., 2015; Oguz et al., 2014) demonstrated that residual motion effects can lead to systematic errors in estimation of WM FA. Furthermore, age-related differences in participant motion have been shown to obscure observed developmental changes in WM microstructure (Roalf et al., 2016). Participants from clinical populations may also be more likely than healthy controls to exhibit head motion during DWI acquisition, resulting in spurious group differences in diffusion scalar measures that can be attenuated by including head motion as a nuisance regressor (Yendiki et al., 2014). Although the impact of head motion on diffusion scalar metrics has been well-characterized in previous work, the downstream effects of motion on network-based measures of structural connectivity have not been systematically examined.

Here, we leveraged DTI data collected as part of the Philadelphia Neurodevelopmental Cohort (PNC), a large population-based study of human brain development (Satterthwaite et al., 2014, 2016), to evaluate the impact of participant motion on structural connectivity. We hypothesized that subtle variation in head motion would systematically bias estimates of structural connectivity and confound inferences regarding brain development. Since head motion can result in both the overestimation *and* underestimation of diffusion anisotropy depending on regional FA and SNR (Farrell et al., 2007; Jones and Basser, 2004; Landman et al., 2008; Tijssen et al., 2009), participant motion could promote spurious streamline propagation in low-FA regions and premature streamline termination in high-FA regions. Moreover, we expected that motion would have a differential impact on structural connectivity depending on specific attributes of each network edge. Specifically, we predicted that motion would inflate estimates of structural connectivity for potentially spurious, low-FA connections that were primarily short-range, while simultaneously diminishing estimates of structural connectivity for long-range, high-FA connections that were consistently reconstructed across participants. To test these hypotheses, structural connectivity was measured in 949 youth (ages 8-23 years old) after constructing brain networks using both deterministic and probabilistic tractography.

## MATERIALS AND METHODS

### Participants and data acquisition

The DTI datasets used in this study (*N*=949) were collected as part of the Philadelphia Neurodevelopmental Cohort (PNC; Satterthwaite et al., 2014, 2016) and selected on the basis of health and data quality criteria. All participants included in this study were ages 8-23 years old at the time of scan, lacked gross structural brain abnormalities (Gur et al., 2013), were free from medical conditions that could impact brain function (Merikangas et al., 2010), were not taking psychotropic medication at the time of the scan, and passed a rigorous manual quality insurance protocol involving visual inspection of all 71 volumes (Roalf et al., 2016). The exclusion of participants with gross artifact due to head motion, eddy currents, susceptibility artifacts, and/or other scanner artifacts allowed us to more rigorously evaluate the impact of subtle in-scanner motion on estimates of structural connectivity (for further details regarding manual quality assurance, see below).

### Image acquisition

Structural and diffusion MRI scans were acquired using the same 3T Siemens Tim Trio whole-body scanner and 32-channel head coil at the Hospital of the University of Pennsylvania. DTI scans were acquired using a twice-refocused spin-echo (TRSE) single-shot echo-planar imaging (EPI) sequence (TR = 8100ms, TE = 82ms, FOV = 240mm / 240mm; Matrix = RL: 128, AP:128, Slices:70, in-plane resolution (x and y) 1.875 mm; slice thickness = 2mm, gap = 0; flip angle = 90°/180°/180°, volumes = 71, GRAPPA factor = 3, bandwidth = 2170 Hz/pixel, PE direction = AP). This sequence used a four-lobed diffusion encoding gradient scheme combined with a 90-180-180 spin-echo sequence designed to minimize eddy-current artifacts. For DTI acquisition, a 64-direction set was divided into two independent 32-direction imaging runs in order to increase the likelihood of scan completion for young subjects. Each 32-direction sub-set was chosen to be maximally independent such that they separately sampled the surface of a sphere (Jones et al., 2002). The complete sequence was approximately 11 minutes long, and consisted of 64 diffusion-weighted directions with b=1000s/mm^2^ and 7 interspersed scans where *b*=0 s/mm^2^. The imaging volume was prescribed in axial orientation covering the entire cerebrum with the topmost slice just superior to the apex of the brain (Satterthwaite et al., 2014). In addition to the DTI scan, a map of the main magnetic field (i.e., B0) was derived from a double-echo, gradient-recalled echo (GRE) sequence, allowing us to estimate field distortions in each dataset.

### Structural image processing and quality assurance

High-resolution structural images were processed using FreeSurfer (version 5.3) (Fischl, 2012), and cortical and subcortical gray matter was parcellated according to the Lausanne atlas (Cammoun et al., 2012), which includes a 233-region subdivision of the Desikan-Killany anatomical atlas (Desikan et al., 2006). Parcellations were defined in native structural space and co-registered to the first *b*=0 volume of each participant’s diffusion image using boundary-based registration (Greve and Fischl, 2009). All participants included in this study passed quality assurance procedures for the raw T1 input image and following FreeSurfer reconstruction (Rosen et al., 2017).

### DTI preprocessing

The two consecutive 32-direction acquisitions were merged into a single 64-direction time-series. Skull-stripping was performed by registering a binary mask of a standard fractional anisotropy (FA) map (FMRIB58 FA) to each subject’s DTI image using FLIRT (Jenkinson et al., 2002). Eddy currents and subject motion were estimated and corrected using the FSL *eddy* tool (Andersson and Sotiropoulos, 2016). This procedure uses a Gaussian Process to simultaneously model the effects of eddy currents and head motion on diffusion-weighted volumes, resampling the data only once. Diffusion gradient vectors were then rotated to adjust for subject motion estimated by *eddy* (Leemans and Jones, 2009). After the field map was estimated, distortion correction was applied to DTI images using FSL’s FUGUE (Jenkinson et al., 2012).

### Manual DTI quality assurance

Manual quality assurance for the DTI images was performed prior to diffusion model fitting, tractography, and structural brain network construction. Specifically, each volume of the acquisition (*n*=71) was evaluated for the presence of artifact, and the total number of impacted volumes over the whole series was recorded (Roalf et al., 2016). This scoring was based on previous work characterizing the detrimental impact of removing diffusion-weighted volumes when estimating the diffusion tensor (Chen et al., 2015; Jones and Basser, 2004). Data was defined as “Poor” if more than 14 (20%) volumes contained artifact, “Good” if it contained 1-14 volumes with artifact, and “Excellent” if no visible artifacts were detected in any volumes. All 949 participants included in the present study had DTI datasets identified as “Good” or “Excellent”. While including participants with poor data quality would undoubtedly lead to larger observed motion effects, in this study we sought to characterize the impact of subtle in-scanner motion in a sample that would typically be included in studies of brain development.

### Diffusion model fitting, tractography, and brain network construction

#### Probabilistic Pipeline

A ball-and-sticks diffusion model was fitted to each subject’s DTI data using FSL *bedpostx*, which uses Markov chain Monte Carlo sampling to build distributions on principal fiber orientation and diffusion parameters at each voxel (Behrens et al., 2007). In contrast to tensor-based approaches, this allowed us to model up to two crossing fibers per voxel, enhancing sensitivity to more complex white matter architecture. Probabilistic tractography was run using FSL *probtrackx*, which repetitively samples voxel-wise fiber orientation distributions to model the spatial trajectory and strength of anatomical connectivity between specified seed and target regions (Behrens et al., 2007). Here, we defined seeds in native T1 space by dilating the original 233-region gray matter parcellation by 2mm and then masking dilated regions by the boundary of each subject’s white matter (WM) segmentation. Once defined for each subject, the seed mask was co-registered to the first *b* = 0 volume of each subject’s diffusion image using boundary-based registration (Greve and Fischl, 2009).

Each cortical and subcortical region defined along the gray-white boundary was selected as a seed region, and its connectivity strength to each of the other 232 regions was calculated using probabilistic tractography. At each seed voxel, 1000 samples were initiated (Baum et al., 2017; Li et al., 2012). We used default tracking parameters (a step-length of 0.5mm, 2000 steps maximum, curvature threshold of 0.02). To increase the biological plausibility of white matter pathways reconstructed with probabilistic tractography, streamlines were terminated if they traveled through the pial surface, and discarded if they traversed cerebro-spinal fluid (CSF) in ventricles or re-entered the seed region (Donahue et al., 2016). This fiber tracking procedure allowed us to construct an undirectional connectivity matrix for each participant, where connection weights were defined as the number of probabilistic streamlines connecting two regions (Donahue et al., 2016; Duarte-Carvajalino et al., 2012; Li et al., 2012). We also calculated alternate connection weights including the mean length of probabilistic streamlines connecting a pair of regions (Donahue et al., 2016), and the connectivity probability - the proportion of streamlines initiated from the seed region that successfully reached the target region (Cao et al., 2013; Johansen-Berg et al., 2005). The procedure for constructing participant connectomes is illustrated in Figure 1.

**Figure 1.**
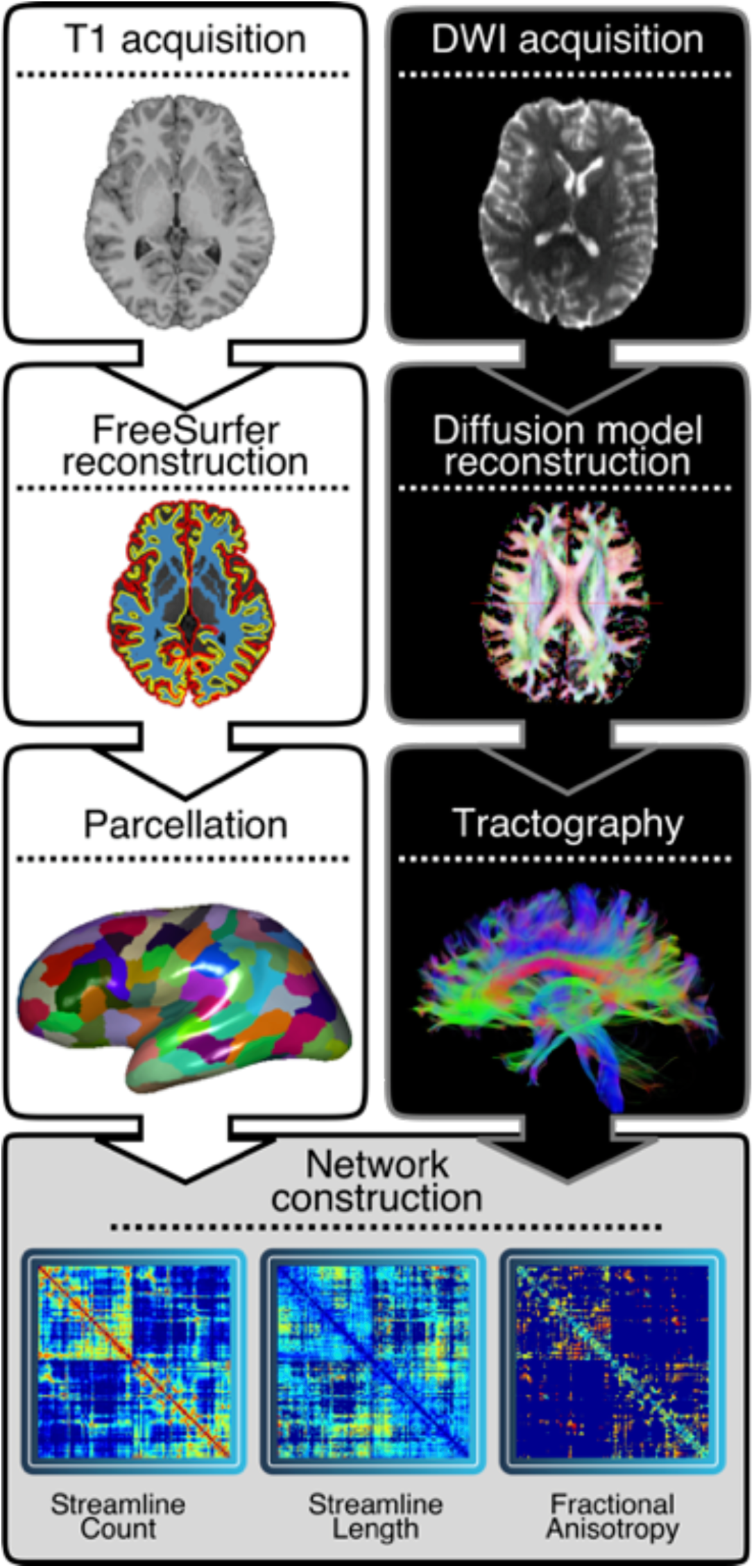
Connectome construction. For each subject (*n*=949, ages 8-23 years), the T1 image was processed using FreeSurfer and parcellated into 233 cortical and subcortical network nodes on a subject-specific basis. A ball-and-stick diffusion model was fit to each subject’s DTI data and probabilistic tractography was run with FSL *probtrackx*, initiating 1,000 streamlines in each seed voxel identified at the gray-white boundary for each node. Edge weights in 233×233 symmetric connectivity matrices derived from probabilistic tractography were defined by the number of streamlines connecting a node pair, and by the mean length of streamlines connecting a node pair. Alternatively, the diffusion tensor was fit to the DTI data and deterministic streamline tractography was used to create a symmetric connectivity matrix (233×233), where the primary edge weight was defined by calculating the mean fractional anisotropy (FA) along streamlines connecting a node pair.

#### Deterministic Pipeline

DTI data was imported into DSI Studio software and the diffusion tensor was estimated at each voxel (Yeh et al., 2013). Whole-brain fiber tracking was run for each subject in DSI Studio using a modified fiber assessment by continuous tracking (FACT) algorithm with Euler interpolation. Network nodes were defined by dilating the 233-region gray matter parcellation by 4mm to extend labels beyond the gray-white boundary to include deep white matter (Baum et al., 2017; Gu et al., 2015). Following standard procedures, we used whole-brain tractography to initiate 1,000,000 streamlines while removing all streamlines with length less than 10mm or greater than 400mm. Fiber tracking was performed with an angular threshold of 45°, a step size of 0.9375mm, and a fractional anisotropy (FA) threshold determined empirically by Otzu’s method, which optimizes the contrast between foreground and background (Yeh et al., 2013). As in previous studies of human structural brain networks, connection weights were defined by calculating the average FA along each streamline connecting a node pair (Baum et al., 2017; Bohlken et al., 2016; Mišić et al., 2016; van den Heuvel and Sporns, 2011). This measure of connection strength is thought to reflect underlying microstructural properties of WM such as myelination or axonal density (Chang et al., 2017; Gulani et al., 2001; Paus, 2010; Takahashi et al., 2002). To evaluate motion effects on the distance of reconstructed fiber pathways, we also defined connection weights as the mean length of streamlines connecting a node pair. Supplementary analyses evaluated motion effects on structural connectivity when edge weights were defined by the average inverse MD along streamlines connecting a node pair (Friedrichs-Maeder et al., 2017; Hagmann et al., 2010; Wierenga et al., 2016), and by the deterministic streamline count (Bassett et al., 2011; van den Heuvel et al., 2015).

### Quantifying in-scanner head motion during DTI acquisition

In-scanner head motion was primarily measured by the mean relative volume-to-volume displacement between the higher SNR *b*=0 images (*n*=7), which summarizes the total translation and rotation in 3-dimensional Euclidean space (Roalf et al., 2016; Satterthwaite et al., 2012; Van Dijk et al., 2012). To determine the specificity of our results, we also conducted supplementary analyses to evaluate whether alternative measures of head motion and data quality impacted structural connectivity. These measures included the following: (*1*) average volume-to-volume translation, (*2*) average volume-to-volume rotation calculated across all 71 volumes (Yendiki et al., 2014), (*3*) mean voxel outlier count, and (*4*) average temporal signal-to-noise ratio (TSNR) defined using the 64 diffusion-weighted volumes, as described in detail in Roalf et al. (2016).

### Edge consistency

Deterministic and probabilistic tractography algorithms for reconstructing WM connectivity face a well-characterized tradeoff between connectome specificity and sensitivity (Knösche et al., 2015; Thomas et al., 2014; Zalesky et al., 2016). Thus, identifying and controlling for the influence of false positives and false negatives remains a critical issue in connectome construction, as both the failure to reconstruct “real” connections and the inclusion of spurious connections can substantially bias group-level inferences on network organization (Drakesmith et al., 2015; Zalesky et al., 2016). Prior work has demonstrated how partial volume effects and complex WM geometry can result in premature streamline termination during tractography when termination criteria are based on WM curvature and anisotropy thresholds (Smith 2012, 2013; Girard 2014; Vos 2011). Notably, head motion can artificially inflate FA estimates in low anisotropy regions and reduce FA in highly coherent WM regions (Farrell et al., 2007; Jones and Basser, 2004; Landman et al., 2008; Ling et al., 2012; Tijssen et al., 2009), potentially compounding these tractography biases by promoting spurious streamline propagation in low-FA regions and premature streamline termination in high-FA regions. Moreover, we sought to delineate whether head motion differentially impacted structural connectivity depending on the inter-subject consistency of edge reconstruction.

For dense brain networks derived from probabilistic tractography (mean density= 70.62%, SD= 7.36%), edge consistency was defined by the coefficient of variation for each edge weight across subjects (Roberts et al., 2017). As in prior work, for relatively sparse brain networks derived from deterministic tractography (mean density=13.95%, SD=0.9%), edge consistency was defined by the percentage of subjects with a non-zero weight for a given edge (de Reus and van den Heuvel, 2013).

### Statistical analysis: group-level motion effects

The effect of in-scanner head motion on structural connectivity was estimated using a partial correlation for each network edge while controlling for potentially confounding demographic variables (age, age^2^, and sex). To assess the significance of the third level correlation between edge-level motion effects (partial *r* coefficients) and edge consistency, we performed an edge-based permutation test. Specifically, we recalculated the correlation between edge-level motion effects and edge consistency after permuting edge consistency 10,000 times. Then, we determined where the observed correlation between motion effects and edge consistency fell relative to this null distribution. In light of prior work characterizing distance-dependent motion effects on functional connectivity (Ciric et al., 2017; Power et al., 2012; Satterthwaite et al., 2012), this permutation procedure was repeated to assess the third-level correlation between motion effects and mean streamline length.

### Consistency-based thresholding

After evaluation of the relationship between in-scanner motion and structural connectivity, we next evaluated the impact of thresholding procedures on such effects. Thresholding approaches are commonly applied to human brain networks in order to reduce the prevalence of spurious false positive connections that may bias group-level inferences on brain network topology (Drakesmith et al., 2015; Roberts et al., 2017; Rubinov and Sporns, 2010; Zalesky et al., 2016). While one common thresholding approach involves removing a subset of the weakest edges in a group-average connection matrix (Rubinov and Sporns, 2010), this approach often results in the elimination of relatively weak, long-range connections that may be particularly important for global network topology (Roberts et al., 2017; van den Heuvel et al., 2012). In contrast, consistency-based thresholds retain both short-and long-range connections that are consistently reconstructed across subjects (Roberts et al., 2017). In the present study, we sought to delineate motion effects on structural connectivity after eliminating potentially spurious network edges. To this end, we applied consistency-based thresholds to brain networks derived from both probabilistic (Roberts et al., 2017) and deterministic tractography (de Reus and van den Heuvel, 2013).

For networks derived from probabilistic tractography, we evaluated motion effects on edge strength, node strength, and total network strength across ten consistency-based thresholds (0-90^th^ percentile probabilistic edge consistency). In agreement with previous studies using deterministic tractography, which have applied group-level thresholds based on the percentage of subjects with a given edge rather than percentiles of edge consistency (de Reus and van den Heuvel, 2013; van den Heuvel and Sporns, 2011; Wierenga et al., 2016), we evaluated motion effects on structural connectivity across ten consistency-based thresholds (0-90% deterministic edge consistency). To characterize the severity of motion effects across consistency-based thresholds, we calculated the percentage of network edges and nodes significantly impacted by motion after adjusting for the false discovery rate (FDR; Benjamini and Hochberg, 1995). To assess the stability of motion effects across consistency-based thresholds, we generated 100 bootstrap samples defined using 80% of the dataset (*N*=760). The percentage of edges significantly impacted by motion and the effect of motion on total network strength were calculated across consistency-based thresholds for each bootstrap sample.

### Statistical analysis: group-level age effects and mediation analysis

As a final step, we examined whether observed age effects on structural connectivity were mediated by age-related differences in head motion. Sobel tests were performed for each network edge exhibiting significant age effects following FDR correction (Sobel, 1982). Specifically, for the subset of edges where age-related differences in head motion significantly mediated observed age effects on structural connectivity, we performed 10,000 permutations of an edge-level index defining mediation effects as “positive” or “negative” depending on the value of the Sobel *Z* statistic. For each permutation, we calculated the difference in mean edge consistency between the randomly labeled “positive” and “negative” mediation effects, and ultimately compared the observed difference in mean edge consistency to this null distribution.

## RESULTS

### In-scanner head motion systemically impacts estimates of structural connectivity in a consistency-dependent manner

When edge weights were defined by the number of probabilistic streamlines connecting a node pair, 12.12% of all network edges were significantly impacted by motion (Figure 2A). Notably, both positive and negative motion effects were observed: motion diminished the strength of 56.73% of these edges and increased the strength of the remaining 43.27%. We found that the direction and strength of motion effects on streamline count were correlated with edge consistency (*r*=-0.35, permuted *p* < 0.0001; Figure 2B) as well as with mean streamline length (*r*=-0.21, permuted *p* < 0.0001). When edge weights were defined by the mean length of probabilistic streamlines connecting a node pair, 62.38% of all network edges were significantly impacted by motion (Figure 2C). Specifically, motion diminished streamline length in nearly all (98.32%) of the impacted edges, supporting the view that motion artifact might increase the likelihood of premature streamline termination. As for streamline count, the direction and strength of motion effects on streamline length were correlated with edge consistency (*r*=-0.59, permuted *p* < 0.0001; Figure 2D) and with mean streamline length (*r*=-0.73, permuted *p* < 0.0001).

**Figure 2.**
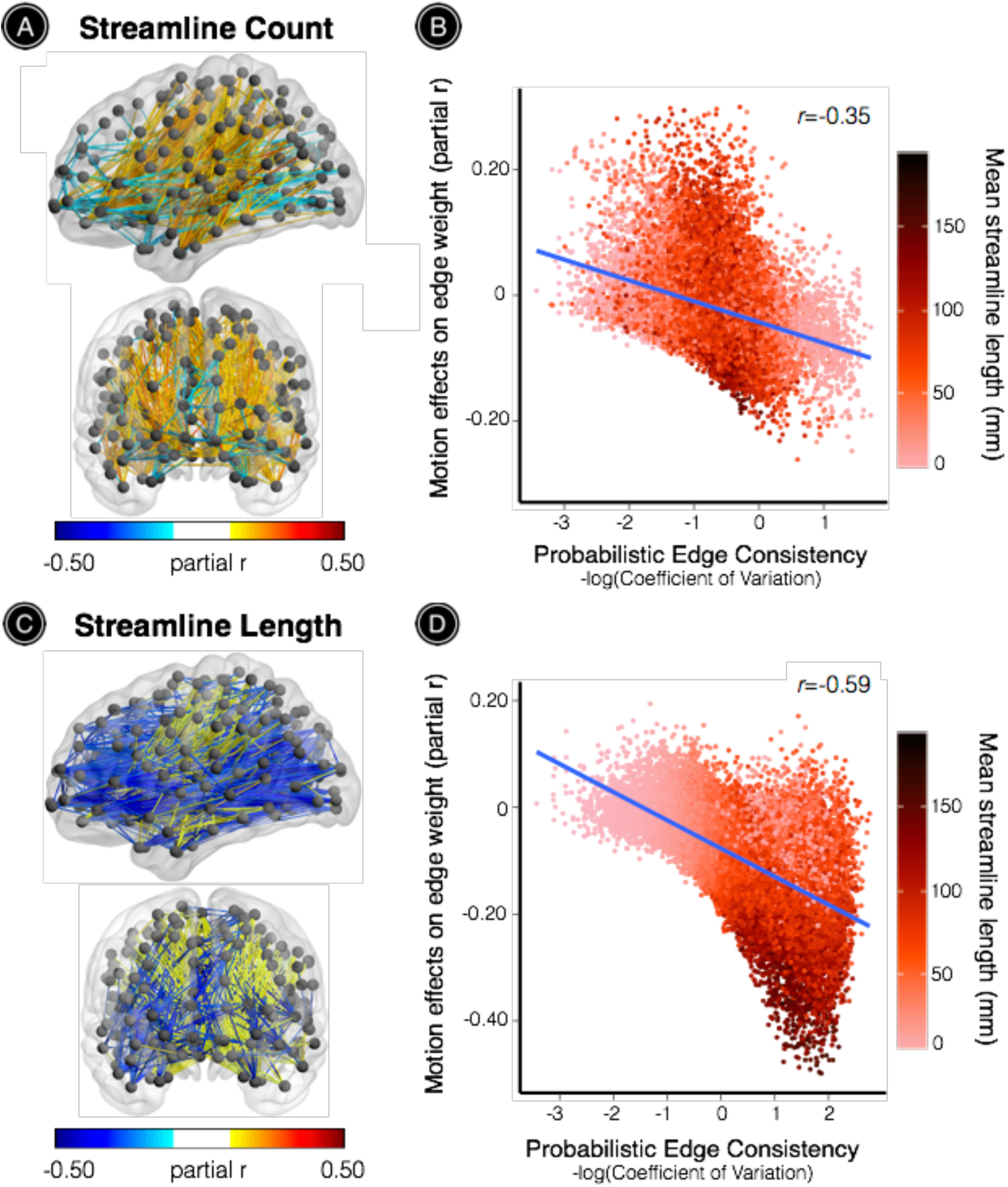
Motion effects on structural connectivity are driven by edge consistency and streamline length. The effect of in-scanner head motion on structural connectivity was estimated using a partial correlation for each network edge while controlling for age, age^2^, and sex. (**A**) When edge weights were defined by the number of probabilistic streamlines connecting a node pair, 12.12% of all network edges were significantly impacted by motion (56.73% negative effects). (**B**) The direction and strength of motion effects were significantly correlated with edge consistency (*r*=-0.35) and with mean streamline length (*r*=-0.21). (**C**) When edge weights were defined by the mean length of probabilistic streamlines connecting a node pair, 62.38% of all network edges were significantly impacted by motion (98.32% negative effects). (**D**) The strength and direction of motion effects were significantly correlated with edge consistency (*r*=-0.59) and mean streamline length (*r*=-0.73). All statistical inferences were adjusted for multiple comparisons using FDR (*Q* < 0.05). The significance of all third-level correlations was evaluated using 10,000 permutations (permutation-based *p* < 0.0001).

We also evaluated the impact of head motion on structural connectivity using brain networks derived from deterministic tractography. When edge weights were defined by the mean FA along deterministic streamlines connecting a node pair, 13.72% of all network edges were significantly impacted by motion (Figure 3A). Specifically, motion diminished the strength of 45.94% of these edges and increased the strength of the remaining 54.06%. As for probabilistic tractography, the impact of motion was dependent on both consistency and streamline length: the direction and strength of motion effects were correlated with edge consistency *(r=-0.50*, permuted *p* < 0.0001; Figure 3B) and with mean streamline length *(r=-0.48*, permuted *p* < 0.0001). When edge weights were defined by the length of deterministic streamlines connecting a node pair, 10.17% of all network edges were significantly impacted by motion (Figure 3C). Motion diminished the strength of 35.5% of these edges and enhanced the strength of the remaining 64.65%. The direction and strength of motion effects were correlated with edge consistency *(r=-0.34*, permuted *p* < 0.0001; Figure 3D) and with mean FA *(r=-0.39*, permuted *p* < 0.0001).

**Figure 3.**
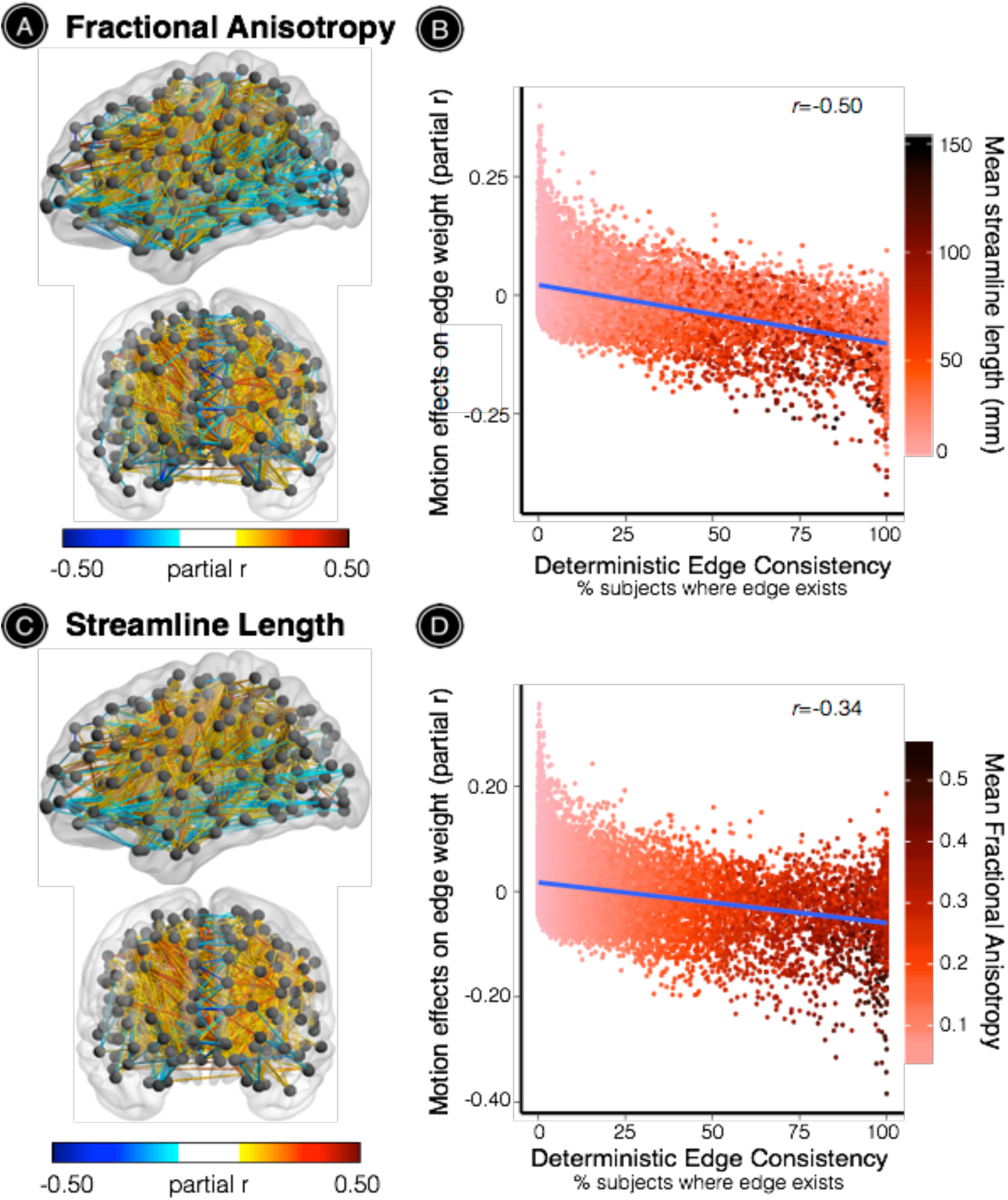
Consistency-and length-driven motion effects are similar using deterministic tractography. (**A**) When edge weights were defined by the average FA along deterministic streamlines connecting a node pair, 13.72% of all network edges were significantly impacted by motion (54.06% positive effects). (**B**) The direction and strength of motion effects were significantly associated with edge consistency (*r*=-0.50) and with mean streamline length (*r*=-0.50). (**C**) When edge weights were defined by the mean length of streamlines connecting a node pair, 10.17% of all network edges were significantly impacted by motion (64.65% positive effects). (**D**) The direction and strength of motion effects were significantly associated with edge consistency (*r*=-0.34) and with mean FA (*r*=-0.39). All statistical inferences were adjusted for multiple comparisons using FDR (*Q* < 0.05). The significance of all third-level correlations was evaluated using 10,000 permutations (permutation-based *p* < 0.0001).

To further disentangle the associations between edge-level motion effects, edge consistency, and streamline length, we plotted these relationships for the subset of edges significantly impacted by motion (FDR *Q*<0.05). For brain networks derived from probabilistic tractography, we observed a quadratic relationship between streamline length and edge consistency. Head motion significantly enhanced the strength of relatively short-range, low-consistency network edges, and diminished the strength of high-consistency network edges, which included both short-and long-range connections (Figure 4A). Similarly, for brain networks derived from deterministic tractography, head motion significantly enhanced FA along relatively short-range, low-consistency network edges, and diminished FA along relatively long-range, high-consistency network edges (Figure 4B).

**Figure 4.**
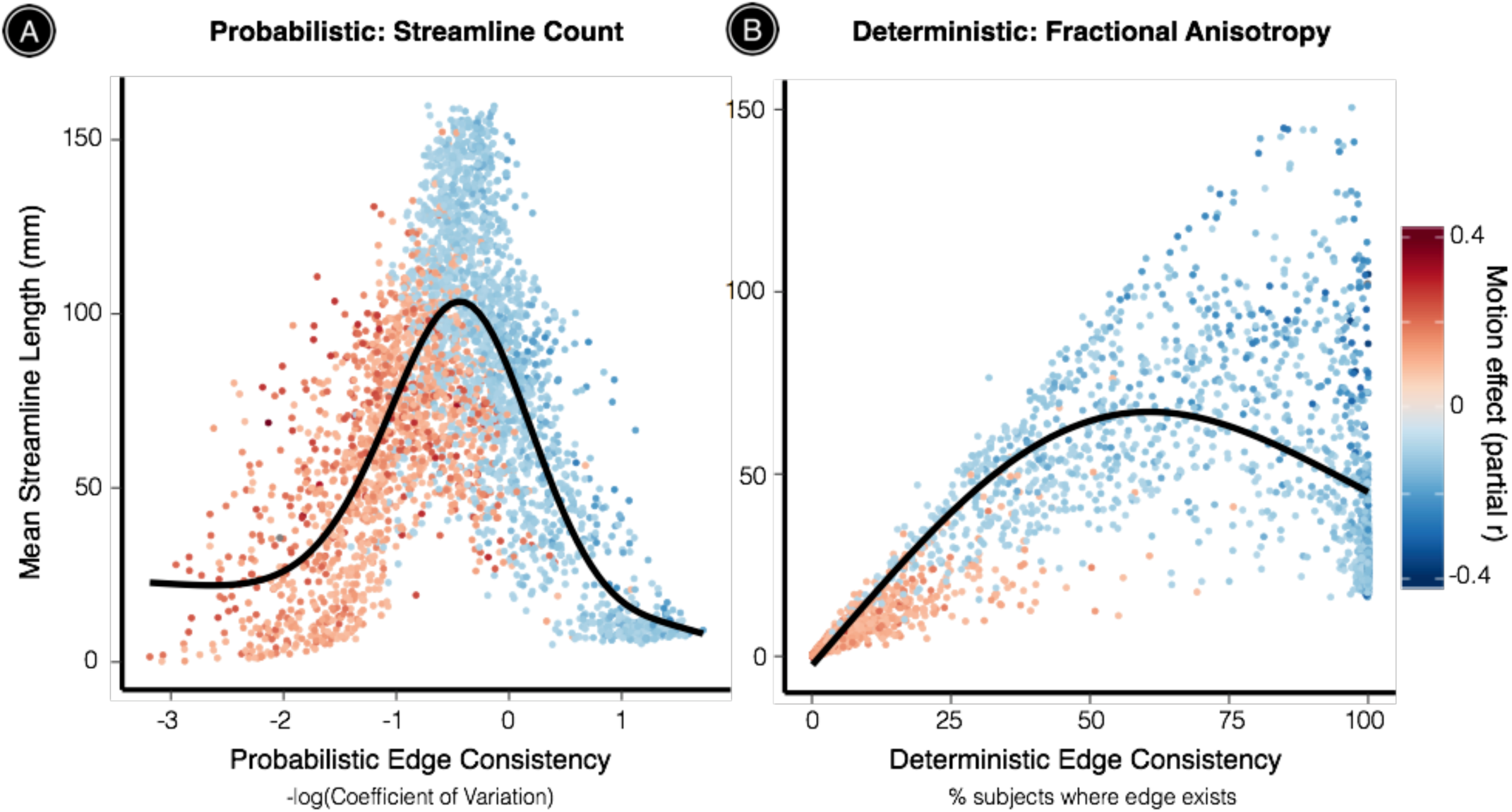
Head motion enhances connectivity in short-range, low-consistency edges, and diminishes connectivity in high-consistency edges. (**A**) In networks defined using probabilistic tractography, edge consistency exhibited a quadratic relationship with mean streamline length. Notably, head motion significantly enhanced the strength of relatively short-range, low-consistency network edges. Further, head motion diminished the strength of relatively high-consistency network edges, which included both short-and long-range connections. (**B**) For deterministic tractography, edge consistency exhibited a parabolic relationship with mean streamline length. In agreement with results from probabilistic tractography, head motion significantly enhanced the strength of relatively short-range, low-consistency network edges. Further, head motion diminished the strength of relatively long-range, high-consistency network edges. Black line represents the best fit from a general additive model with a penalized spline. Data shown for edges with significant motion effects (FDR *Q* < 0.05).

Notably, we observed highly consistent motion effects on structural connectivity when using a variety of other edge weight definitions for networks derived from both deterministic and probabilistic tractography (Supplemental Figure 1), including connectivity probability, inverse MD, and deterministic streamline count. We also demonstrated that alternative measures of data quality, such as the mean framewise translation and rotation, the number of of mean voxel intensity outliers across diffusion-weighted volumes, and TSNR, all exhibited similar effects on structural connectivity (Supplemental Figure 2).

### Motion effects are exacerbated across consistency-based thresholds

We applied ten consistency-based thresholds to networks derived from both probabilistic and deterministic tractography in order to evaluate the impact of head motion on edge strength, node strength, and total network strength after eliminating potentially spurious network edges. For networks derived from probabilistic tractography, the percentage of edges significantly impacted by head motion increased monotonically across consistency-based thresholds ranging from (12.12 - 32.3%; Figure 5A). Motion had a profound impact on network properties at the nodal level, significantly diminishing the strength of 82.4-89.7% nodes across consistency-based thresholds. After retaining only the top 50^th^ percentile of edges based on intersubject consistency, head motion had a significant negative effect on the strength of 83.69% nodes, with particularly strong effects observed in middle frontal gyrus, precuneus, and cingulate cortex (Figure 5B). Total network strength was also significantly diminished by head motion across all consistency-based thresholds (partial *r* ranged between −0.298 and −0.314; Figure 5C).

**Figure 5.**
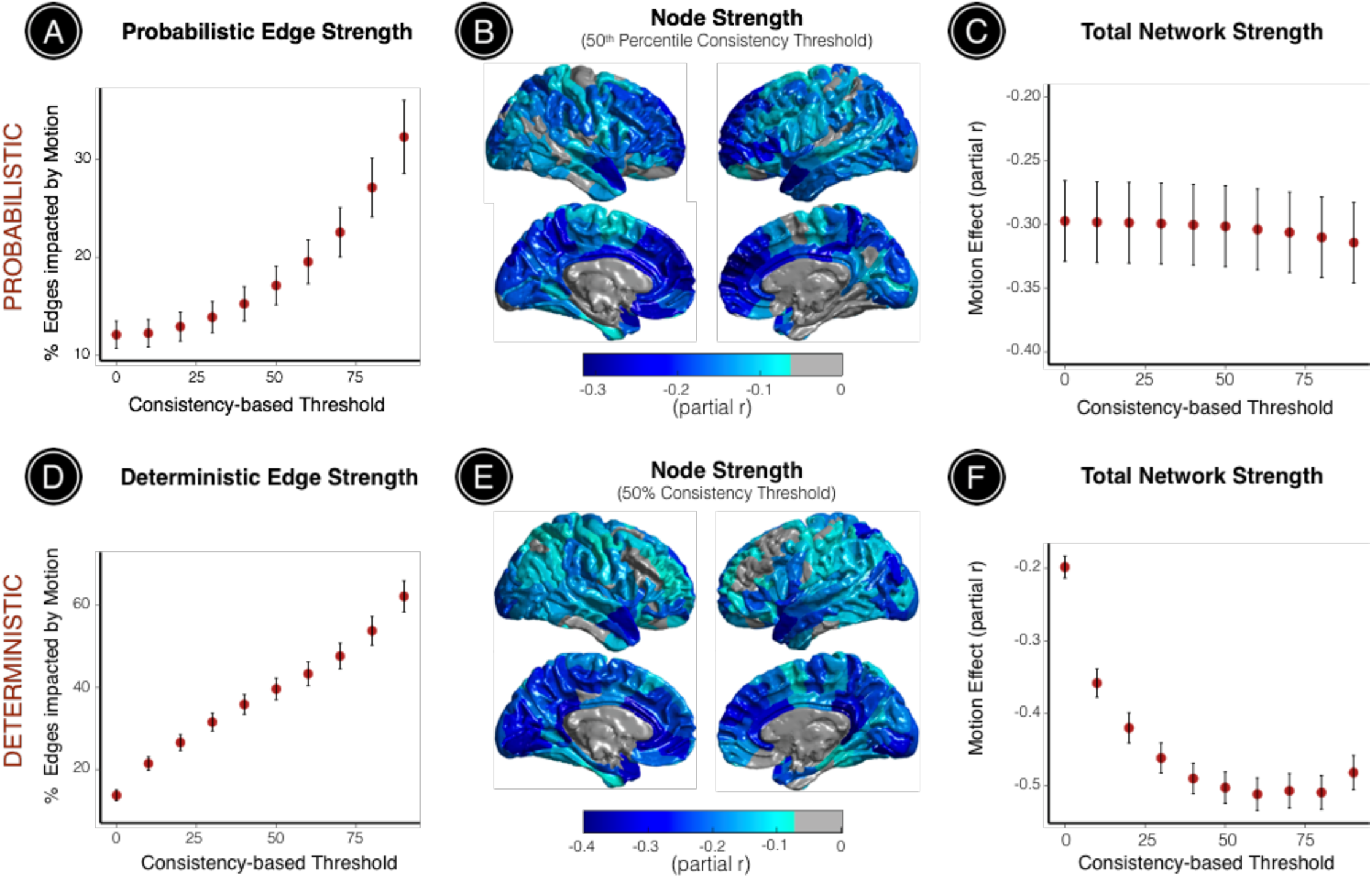
Head motion systematically impacts structural connectivity across consistency-based thresholds at the level of network edges, nodes, and total network strength. Motion effects on probabilistic edge strength, node strength, and total network strength were assessed across a range of consistency-based thresholds (ten thresholds, 0-90^th^ percentile consistency). (**A**) The percentage of edges significantly impacted by head motion increased monotonically across consistency-based thresholds (12.12 −32.3%). (**B**) After eliminating all edges with consistency below the 50^th^ percentile, head motion significantly diminished the strength of 83.69% nodes, with particularly strong effects observed in middle frontal gyrus, precuneus, and cingulate cortex. (**C**) While the effect was stable across consistency-based thresholds, head motion significantly diminished total network strength at each threshold. Motion effects on deterministic edge strength, node strength, and total network strength were assessed across ten consistency-based thresholds (0-90% deterministic edge consistency). (**D**) The percentage of deterministic network edges significantly impacted by head motion increased monotonically across consistency-based thresholds (13.72 - 62.19%). (**E**) After eliminating edges that existed in less than 50% of participant connection matrices, head motion significantly diminished the strength of 89.27% nodes, with particularly strong effects observed in the precuneus and medial brain regions including the anterior and posterior cingulate. (**F**) Head motion also significantly diminished total network strength across all consistency-based thresholds, particularly at more stringent thresholds. These results suggest that global strength normalization approaches may be confounded by individual differences in head motion during acquisition. All statistical inferences were adjusted for multiple comparisons using FDR (*Q* < 0.05). Black bars correspond to the standard deviation of 100 bootstrapped samples encompassing 80% of the dataset (*N*=760).

For networks derived from deterministic tractography, the percentage of network edges significantly impacted by head motion also increased monotonically across consistency-based thresholds (13.72 - 62.19%; Figure 5D). Motion also reduced the strength of a large percentage of network nodes (43.35 - 98.71%). After retaining only edges that were reconstructed in more than 50% of participant connection matrices, head motion significantly reduced the strength of 89.27% nodes, with particularly strong effects observed in the precuneus and medial brain regions including the anterior and posterior cingulate cortex (Figure 5E). Head motion significantly reduced total network strength across all consistency-based thresholds, with stronger effects observed at more stringent thresholds (partial *r* varied between −0.20 and −0.51; Figure 5F). These results demonstrate the impact of motion artifact on structural connectivity across topological scales, thresholding procedures, and network construction methods.

### Age effects on structural connectivity are inflated and obscured by head motion

As a final step, we evaluated whether motion could systematically bias estimates of structural network development during youth. Even in our sample of 949 youths with high-quality, low-motion DTI data (mean participant motion=0.46mm, SD=0.41mm), head motion was negatively correlated with age such that younger participants tended to move significantly more than older participants (*r*=-0.17, *p*=3.01 × 10^−7^; Figure 6A). While controlling for participant sex, significant age effects were observed in 25.63% of probabilistic network edges and 6.87% of deterministic network edges for unthresholded networks. We tested whether these significant age effects were mediated by participant motion using the Sobel test. Figure 6B illustrates that positive Sobel *Z* values can reflect either *inflated* positive age effects or *obscured* negative age effects, where in both cases motion decreases the strength of network edges that undergo significant age-related change. Similarly, negative Sobel Z values can reflect either *inflated* negative age effects or *obscured* positive age effects, where in both cases motion increases the strength of network edges that undergo significant age-related change. For brain networks derived from probabilistic tractography, 7.42% of edges with observed age effects were significantly mediated by age-related differences in head motion (39.34% positive mediation, 60.66% negative mediation; Figure 6C). Notably, network edges with significant positive mediation effects had higher edge consistency compared to connections with significant negative mediation effects (permutation-based *p* < 0.0001). This result reflects the fact that the strength of edges with positive mediation effects are weakened by motion, and negative motion effects are most prominent in high-consistency edges.

**Figure 6.**
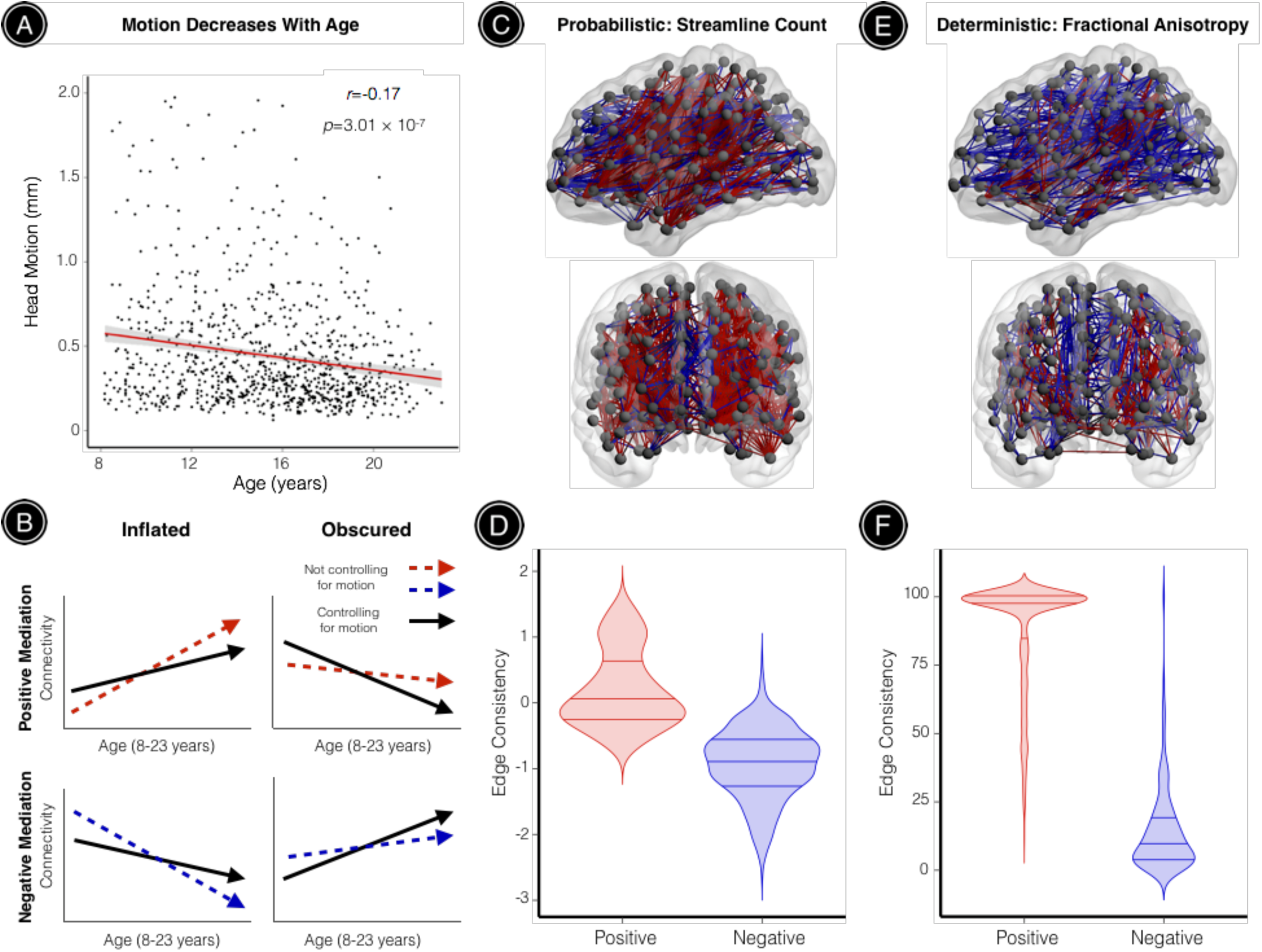
Observed age effects on structural connectivity are both inflated and obscured when age-related differences in head motion are not accounted for. All subjects included in this study passed rigorous manual quality assurance, retaining a sample of relatively high-quality, low-motion DTI datasets. (**A**) Despite this, age-related differences in head motion were still observed: younger participants tended to move significantly more than older participants. (**B**) Mediation analyses across all network edges showing significant age effects demonstrated that observed age effects on structural connectivity were often inflated or obscured when head motion was not accounted for. This schematic illustrates how positive mediation effects can reflect *inflated* positive age effects or *obscured* negative age effects, where in both cases motion decreases the strength of network edges that undergo significant age-related change. Similarly, negative mediation effects can reflect *inflated* negative age effects or *obscured* positive age effects, where in both cases motion increases the strength of network edges that undergo significant age-related change. (**C**) For brain networks derived from probabilistic tractography, significant age effects were observed in 6,927 network edges (25.63%). This visualization highlights 933 (7.42%) of these edges where developmental effects were significantly mediated by age-related differences in head motion. Positive mediation effects were observed for edges where motion significantly reduced connectivity (39.34%), while negative mediation effects were observed for edges where motion significantly increased connectivity (60.66%) (**D**) Network connections exhibiting positive mediation effects had significantly higher edge consistency compared to connections with significant negative mediation effects (permutation-based *p* < 0.0001). (**E**) For brain networks derived from deterministic tractography, significant age effects were observed in 1,578 network edges (6.87%). This visualization highlights 810 (51.33%) of these edges where developmental effects were significantly mediated by age-related differences in head motion. Again, both positive (81.36%) and negative (18.64%) mediation effects were observed. (**F**) As seen in the probabilistic data, network connections with significant positive mediation effects had significantly higher edge consistency compared to connections with significant negative mediation effects (permutation-based *p* < 0.0001). Red connections represent significant positive mediation results; blue connections represent significant negative mediation results.

For brain networks derived from deterministic tractography, 51.33% of edges with observed age effects were significantly mediated by age-related differences in head motion (87.57% positive mediation, 12.43 % negative mediation; Figure 6E). Consistent with results from probabilistic tractography, network edges with significant positive mediation effects had higher edge consistency compared to connections with significant negative mediation effects (permutation-based *p* < 0.0001; Figure 6F), although this effect was even more pronounced for brain networks derived from deterministic tractography.

## DISCUSSION

Our results demonstrate that subtle variation in participant motion systematically impacts DTI-derived measures of structural connectivity, even following rigorous manual quality assurance. Leveraging DTI data from 949 youths collected as part of the PNC, we found that increased in-scanner head motion was associated with inflated connectivity for low-consistency network edges that were primarily short-range and diminished connectivity for high-consistency edges, which included both long-and short-range connections. Applying group-level thresholds to eliminate potentially spurious connections actually increased the proportion of motion effects on structural connectivity. Furthermore, we demonstrated that age-related differences in head motion could both inflate and obscure developmental inferences on structural connectivity. Our results emphasize that simply applying retrospective motion correction with FSL *eddy* and excluding participants with gross motion artifact are not sufficient for attenuating systematic motion effects on structural connectivity. These findings are particularly important for studies of brain development and neuropsychiatric disorders, where in-scanner motion may be correlated with outcome measures of interest (e.g., participant age, diagnostic group, symptom burden). Together, our results demonstrate that in-scanner micro-movements can have a marked impact on structural connectivity derived from DTI tractography, and they provide a framework for quantifying and controlling for motion-related confounds in studies of structural brain network development.

### Motion effects on structural connectivity are modulated by edge consistency and streamline length

We found that the strength and direction of motion effects on structural connectivity were modulated by inter-subject edge consistency and streamline length. These results are in agreement with studies characterizing the confounding effect of head motion on resting-state functional connectivity (Ciric et al., 2017; Power et al., 2012; Satterthwaite et al., 2012, 2013; Van Dijk et al., 2012; Yan et al., 2013). In diffusion imaging, head motion has been shown to both increase and decrease FA depending on regional tissue anisotropy and signal-to-noise ratio (Aksoy et al., 2008; Farrell et al., 2007; Landman et al., 2008; Ling et al., 2012; Tijssen et al., 2009). Moreover, head motion may bias streamline tractography algorithms that define termination criteria based on voxel-wise FA and step-wise turning angles. Specifically, participant motion may potentially induce a positive FA bias in brain regions with relatively isotropic diffusion, resulting in the spurious propagation of streamlines, while the motion-induced negative FA bias in regions of high anisotropy may result in the premature termination of streamlines. Our results from analyses using streamline length and FA-weighted structural networks further support this premise: head motion increased the length of low-FA, low-consistency connections, and decreased the length of high-FA, high-consistency connections.

When consistency-based thresholds were applied to eliminate potentially spurious network connections as in prior studies (de Reus and van den Heuvel, 2013; Roberts et al., 2017), we found that negative motion effects on edge and node strength became increasingly prevalent. These results are intuitive given that head motion exhibited a particularly strong negative influence on high-consistency network edges. The substantial negative impact of motion on total network strength was stable across all thresholds for networks derived from probabilistic tractography (partial *r* ~ −0.3), and was even more prominent for deterministic networks at more stringent thresholds (partial *r* ~ −0.5). These striking effects on total network strength are particularly notable since many studies assessing intrinsic network topology apply global normalization procedures where each unique edge weight in the individual or group-averaged connectivity matrix is divided by the total network strength (Cao et al., 2013; Dennis et al., 2013; Gong et al., 2009; Li et al., 2012; Yan et al., 2013).

### Age-related differences in head motion both inflate and obscure observed age effects on structural connectivity

The edge consistency-and length-related motion effects on structural connectivity observed in this study have important implications for studies of structural brain network development. While prior work has suggested that short-range WM connections tend to weaken with age while longer-range WM connections become stronger (Collin and van den Heuvel, 2013; Hagmann et al., 2010), our findings suggest that age-related differences in head motion may inflate these age effects in a manner similar to that seen in neurodevelopmental studies of functional connectivity (Fair et al., 2012; Power et al., 2012; Satterthwaite et al., 2012, 2013). Critically, we found that head motion significantly mediated age effects in a consistency-dependent manner, particularly when brain networks were derived from deterministic tractography, where over half of the observed age effects were mediated by motion. Overall, we observed a higher proportion of network edges exhibiting significant age effects using probabilistic tractography, and a smaller proportion of these effects were mediated by age-related differences in motion. Regardless of specific methodological choices during brain network construction, our results demonstrate how subtle differences in participant motion may systematically bias inference regarding the development of structural connectivity in youth.

### Limitations

Several methodological challenges and limitations of the present study should be noted. First, while diffusion tractography methods have been validated using *post-mortem* tract-tracing procedures (Donahue et al., 2016; Knösche et al., 2015; Miranda-Dominguez et al., 2014; van den Heuvel et al., 2015), they remain inherently limited in their ability to fully resolve complex WM trajectories in the human brain, such as fanning and bending fibers (Reveley et al., 2015; Thomas et al., 2014; Zhang et al., 2012). In particular, the relatively low spatial and angular resolution of DTI limits the complexity of diffusion models that can be fitted to the data. State-of-the-art approaches such as neurite orientation dispersion and density imaging (NODDI) leverage multi-shell protocols in combination with high angular resolution diffusion-weighted imaging (HARDI) to enable more nuanced tissue compartment models for assessing WM microstructure and connectivity across the human lifespan (Batalle et al., 2017; Merluzzi et al., 2016; Tuch et al., 2002; Zhang et al., 2012). Critically, tensor-based indices of WM integrity are not sensitive to diffusion within specific intra-voxel tissue compartments, while NODDI can disentangle specific microstructural features such as intra-neurite diffusion (within axons and dendrites), extra-neurite diffusion, and isotropic volume fraction (Zhang et al., 2012). Future studies using NODDI data may help determine whether head motion differentially impacts the diffusion signal in specific tissue compartments.

Second, while a network neuroscience approach provides an attractive way to model pairwise interactions among neural units or brain regions (Bassett and Sporns, 2017), the most optimal method for defining network nodes and edge weights in a biologically meaningful manner remains uncertain (Donahue et al., 2016; Glasser et al., 2016; Gordon et al., 2017; Taylor et al., 2017; Zalesky et al., 2010). Here, we sought to overcome these limitations in part by defining network nodes based on subject-specific neuroanatomical landmarks (Cammoun et al., 2012; Desikan et al., 2006) following rigorous manual and data-driven quality assessments of T1-weighted images (Rosen et al., 2017). Further, our main results were remarkably consistent across a variety of edge weight definitions for networks derived from both deterministic and probabilistic tractography.

Third and finally, we evaluated motion effects on DTI-derived structural connectivity after retrospective correction for field distortions, eddy currents, and participant motion. It should be noted that these preprocessing steps, while commonly applied, can impact diffusion model fitting and tractography results in a non-trivial manner (Alhamud et al., 2015). Future work may benefit from evaluating motion effects on structural connectivity after applying additional procedures for reducing tractography-related biases, such as particle filtering (Girard et al., 2014), anatomically-constrained tractography (ACT) (Smith et al., 2012), or linear fascicle evaluation (LiFE) (Pestilli et al., 2014). Advances in prospective motion correction procedures for diffusion-weighted imaging (Aksoy et al., 2011; Alhamud et al., 2015, 2016) may also help mitigate the impact of motion on structural connectivity.

### Conclusions

In agreement with previous work characterizing motion artifact in structural, functional, and diffusion imaging, we found that in-scanner head motion systematically biases estimates of structural connectivity derived from diffusion tractography and potentially confounds inference on the development of structural brain networks. Based on this data, we recommend that studies of structural brain network topology should quantify data quality, report the relationship between data quality and both subject variables and imaging measures, and control for its influence in analyses through group matching or inclusion of motion as a model covariate. While observed motion effects on structural connectivity were strongest when head motion was measured by the mean relative framewise displacement between interspersed *b*=0 volumes, results suggest that using alternative data quality measures such as nuisance covariates (e.g., outlier count, TSNR) might help to reduce confounding effects in a similar manner when interspersed *b*=0 volumes are not acquired. Taken together, our results delineate the systematic consistency-dependent impact of in-scanner micro-movements on DTI-derived measures of structural connectivity, and emphasize the need for future studies to report and account for the effects of motion artifact.

## ACKNOWLEDGEMENTS

We thank the acquisition and recruitment team, including Karthik Prabhakaran and Jeff Valdez. Thanks to Chad Jackson for data management and systems support and Monica Calkins for phenotyping expertise. Supported by grants from the National Institute of Mental Health: R01MH107703 (TDS), R21MH106799 (DSB & TDS), R01MH112847 (TDS & RTS), and the Lifespan Brain Institute at Penn/CHOP. DSB would like to acknowledge support from the John D. and Catherine T. MacArthur Foundation, the Alfred P. Sloan Foundation, the Army Research Laboratory through contract number W911NF-10-2-0022, the Army Research Office through contract numbers W911NF-14-1-0679 and W911NF-16-1-0474, the National Institute of Health (2-R01-DC-009209-11, 1R01HD086888-01, R01-MH107235, R01-MH107703, R01MH109520, 1R01NS099348 and R21-M MH-106799).The content is solely the responsibility of the authors and does not necessarily represent the official views of any of the funding agencies. The PNC was funded through NIMH RC2 grants MH089983 and MH089924 (REG). Additional support was provided by R01MH107235 (RCG), K01MH102609 (DRR), P50MH096891 (REG), R01NS085211 (RTS), and the Dowshen Program for Neuroscience. Support for developing statistical analyses (RTS & TDS) was provided by a seed grant by the Center for Biomedical Computing and Image Analysis (CBICA) at Penn.

---

**Supplemental Figure 1.**
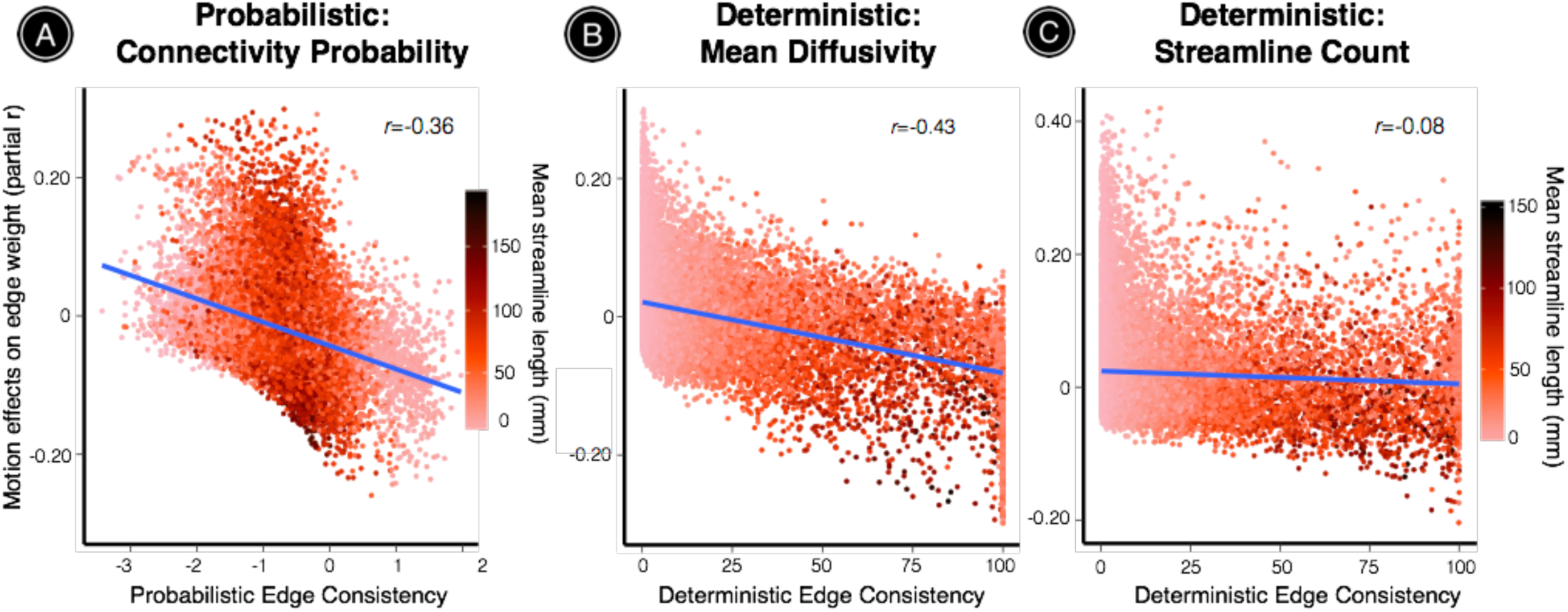
Consistency - and length-driven motion effects using alternative edge weights. (**A**) When edge weights were defined by the connectivity probability between two nodes, 12.66% of all network edges were significantly impacted by motion (57.09% negative effects). The direction and strength of motion effects were significantly associated with probabilistic edge consistency (*r*=-0.36) and mean streamline length (*r*=-0.50). (**B**) When edge weights were defined by the average inverse mean diffusivity (MD) along streamlines connecting a node pair for brain networks derived from deterministic tractography, 13.18% of all network edges were significantly impacted by motion (37.90% negative effects). The direction and strength of motion effects were significantly associated with edge consistency (*r*=-0.43) and with mean streamline length (*r*=-0.43). (**C**) When edge weights were defined by the number of deterministic streamlines connecting a pair of nodes, 12.47% of all network edges were significantly impacted by motion (4.65% negative effects). While the absolute number of edges impacted by motion was highly consistent with that of other edge weights, motion effects on streamline count-weighted networks were only weakly correlated with edge consistency (*r*=-0.08) and with mean streamline length (*r*=-0.17). All statistical inferences were adjusted for multiple comparisons using FDR (*Q* < 0.05). The significance of all third-level correlations was evaluated using 10,000 permutations (permutation-based *p* < 0.0001).

**Supplemental Figure 2.**
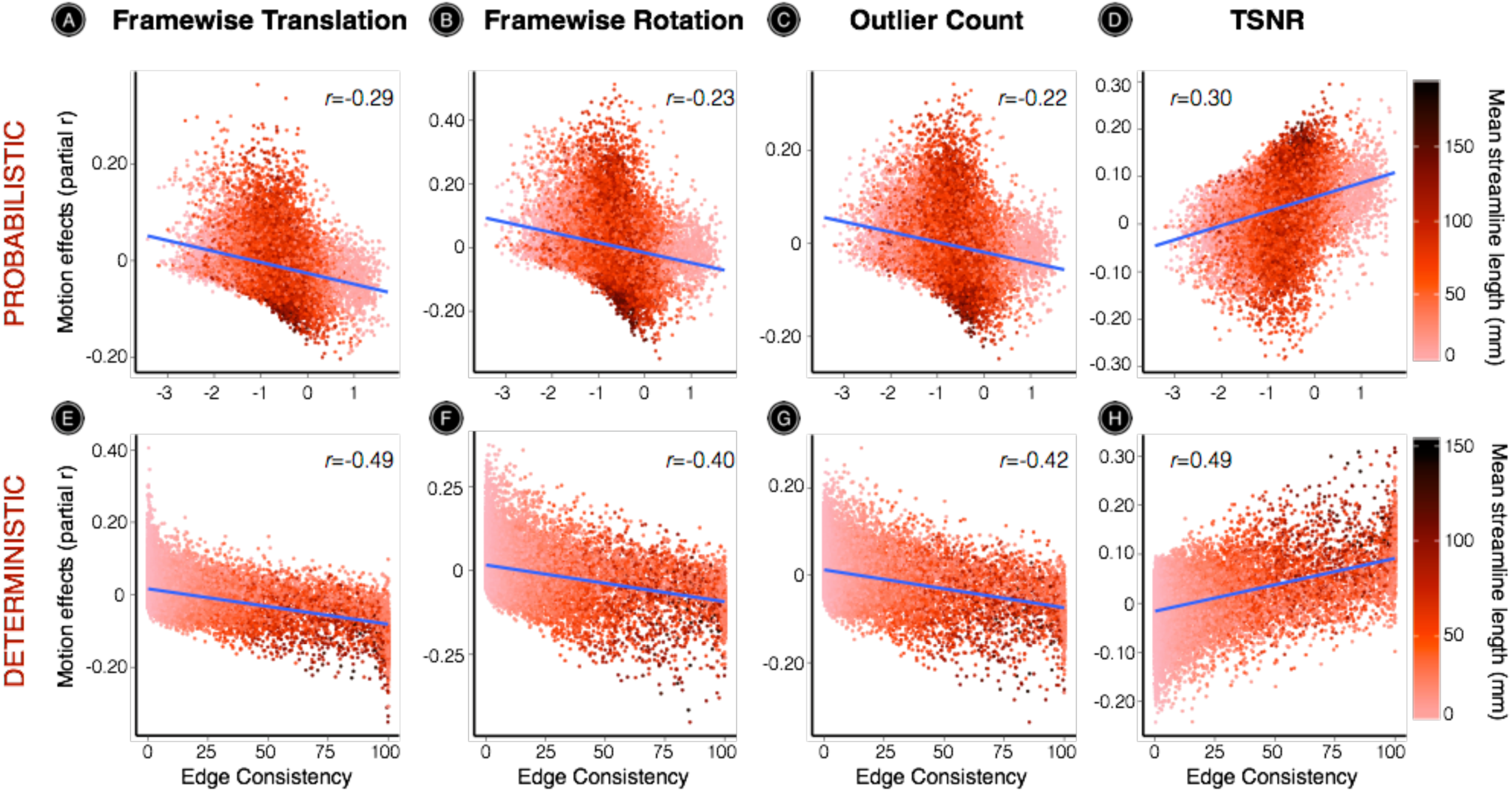
Effects remain highly similar using alternative measures of head motion or DTI data quality. For networks derived from probabilistic tractography, edge weights were defined by the probabilistic streamline count between each pair of nodes. (**A**) Using the translation component of the affine registration from each volume to the first *b*=0 volume, we calculated the average magnitude of translation over all 71 volumes in the scan. This measure of framewise translation significantly impacted the strength of 5.43% of network edges (63.60% positive effects). The direction and strength of these effects were significantly associated with probabilistic edge consistency (*r*=−0.29) and with mean streamline length (*r*=−0.20). (**B**) Using the rotation component of the affine registration from each volume to the first *b*=0 volume, we calculated the average magnitude of rotation over all 71 volumes in the scan. This measure of framewise rotation significantly impacted the strength of 28.41% of network edges (61.36% positive effects). The direction and strength of these effects were significantly associated with probabilistic edge consistency (*r*=−0.23) and mean streamline length (*r*=−0.20). (**C**) The mean voxel outlier count across all 64 diffusion-weighted volumes significantly impacted 13.34% of network edges (61.76% positive effects). The direction and strength of these effects were significantly associated with probabilistic edge consistency (*r*=−0.22) and mean streamline length (*r*=−0.19). (**D**) The mean temporal signal-to-noise ratio (TSNR) across all 64 diffusion-weighted volumes significantly impacted 25.30% of network edges (80.87% positive effects). The direction and strength of these effects were significantly associated with probabilistic edge consistency (*r*=0.30) and mean streamline length (*r*=0.20). For networks derived from deterministic tractography, edge weights were defined by the mean fractional anisotropy (FA) along streamlines connecting each pair of nodes. (**E**) Mean framewise translation significantly impacted the strength of 6.67% of network edges (50.33% positive effects). The direction and strength of these effects were significantly associated with deterministic edge consistency (*r*=−0.49) and mean streamline length (*r*=−0.46). (**F**) Mean framewise rotation significantly impacted the strength of 14.44% of network edges (50.82% positive effects). The direction and strength of these effects were significantly associated with deterministic edge consistency (*r*=−0.40) and mean streamline length (*r*=−0.42). (**G**) The mean voxel outlier count significantly impacted the strength of 7.16% of network edges (46.72% positive effects). The direction and strength of these effects were significantly associated with deterministic edge consistency (*r*=−0.42) and mean streamline length (*r*=−0.42). (**H**) TSNR significantly impacted the strength of 8.92% of network edges (54.81% positive effects). The direction and strength of these effects were significantly associated with deterministic edge consistency (*r*=0.49) and mean streamline length (*r*=0.48). All statistical inferences were adjusted for multiple comparisons using FDR (*Q* < 0.05). The significance of all third-level correlations was evaluated using 10,000 permutations (permutation-based *p* < 0.0001).

